# Assessment of Polarized Piezoelectric SrBi_4_Ti_4_O_15_ Nanoparticles as an alternative antibacterial agent

**DOI:** 10.1101/2021.01.02.425094

**Authors:** Subhasmita Swain, Tapash Ranjan Rautray

## Abstract

Strontium bismuth titanate nanoparticles (SrBi_4_Ti_4_O_15_/SBT NPs,) and their polarized counterparts were prepared to assess their antibacterial efficacy on biomaterials. The structural properties of the SBT NPs were performed by X-ray diffraction and the antibacterial efficacy was evaluated against *Staphyllococcus aureus (S. aureus)* pathogenic bacteria. Significant antibacterial activity of polarized SBT specimen was observed against *S. aureus* bacteria. Results presented in this work confirmed that polarized SBT can effectively combat bacterial growth and prevent biofilm formation activity of pathogenic bacteria and hence they can be used as alternative antimicrobial agents.

## 1. Introduction

Bacterial infection on the biomaterial surfaces has posed a tremendous threat to biomedical implants. The present development and growth in nano material has played a key role in combating bacterial adhesion by showing a deep understanding of the bacteria-implant interaction [1–3]. Bacteria attached to the roughened surfaces in micrometer and even macrometer are protected against abrasion from stress [4]. Moreover, texture of the surface equivalent to the size of bacteria cells is supposed to have higher contact area between the bacteria and the implant thus promoting bacterial adhesion. Subsequently, a present approach for combating bacterial adhesion on the surface of implant is by preparing the surface very smooth. Recently nanomaterials have received tremendous interest because of their effective antibacterial activities [5, 6]. Properties of a material at nanometer scale is significantly different from those of micron sized particles. Nano particles deliver opportunity to enhance the properties and qualities of biomaterials [7, 8]. Nanoparticles have been used as an antibiofilm and antibacterial agent in the growing field of nanotechnology based on drug delivery and gene therapy. The nanoparticles of a particular compound have the ability to enter into cell membrane and interrupt the cell function by killing bacterial cells. However, the bacterial cells sometimes form colonies and get protected by creating biofilms around the colonies which subsequently inhibits antibiotics to act upon them and become resistant [9].

Ferroelectric materials have recently gained significant attention in the fabrication of devices employing non volatile and random access memories [10]. Lead zirconate titanate (PZT) is most wide spread among the ferroelectric materials with a disadvantage of creating environmental pollution because of the presence of lead. Hence lead free environment-friendly materials have constantly been investigated. One among them is strontium-based perovskite that has been found to be highly promising to be used as non-volatile memory applications [11]. SBT has been investigated by several scientists for probable application as piezoelectric biomaterials. SBT has received significant attention because of its anisotropic physical properties, higher Curie temperature, barrier type properties, larger retentivity and lower coercive field [12]. Although numerous methods have been employed in preparing SBT, solid state reaction has been found to be very promising in providing phase pure materials.

Human bone has been found to be a piezoelectric material that can supply the necessary bioelectricity inside the body [13]. It has been affirmed that whenever human bone comes under mechanical stress electric signal is produced that subsequently promotes bone regeneration and growth. Hence, piezoelectric materials are as bone substitutes may be preferred over other biomaterials [14] that can be used as microspheres or scaffolds or bone filler materials. Conversely, SBT has been extensively used as a piezoelectric material because of its piezoelectric coefficient (d33=27 pC/N) and biocompatible properties that is additionally supposed to be an intelligent material for bone regeneration. This property has previously been replicated by other piezoelectric materials such as Barium titanate composites also involving hydroxyapatite [15].

Because of the detrimental effect of bactricidal agents in long term use in human body, their substitute such as polarized biomaterials can be an alternative approach in inhibiting microbial action. Use of external agents including electrical and magnetic inductions on biomaterial surface has been fruitful in reducing bacterial colonies [16]. Although cell behavior is regulated by electric charge inside human body the stimuli generated by the internal electric signal in killing bacterial cells should be widely investigated that can control bacteria activities keeping the human body unharmed [17].

In the present study, effect of piezoelectric materials and their positively polarized counterpart have been employed to study their antibacterial effects on *S. aureus* bacterial cells.

## 2. Material and methods

### 2.1. Synthesis of SBT NPs

Employing solid state reaction method and by using SrCO_3_, Bi_2_O_3_ and TiO_2_, SBT composite was synthesized. Formation of SBT started at 700 °C and its phase pure layer was formed at 900 °C. Elevating the sintering temperature to 1100 °C, SBT started decomposing partially. Thus, to subdue decomposition reaction, SBT composites were cocealed so as to make them highly densified keeping its high purity intact. The phase analysis of the as-formed specimens was carried out using Panalytical X-ray diffractometer (XRD), X’Pert-MPD with PW 3020 vertical goniometer. The data were collected in the range of 20-60° 2θ using CuKα-radiation emitted from an X-ray tube operated at 40 kV and 30 mA [18]. The powdered samples were pelletized using a KBr hydraulic press at 30 Ton pressure to obtain pellets of size 10 mm diameter and 2 mm thickness using Urea binder. The pellets were then sintered at 900 °C for binder burn out.

The polarization of specimens was carried out using a corona poling system (Millman thin films PVT. LTD. Pune, India) at a potential of 2 kV/mm at 550 °C after coating silver paste on both sides of the specimens. The silver coatings were then removed from both sides of pellets and were mirror polished using sand paper. Energy dispersive X-ray fluorescence (EDX) spectroscopic technique was employed to analyze the elemental composition of the as-formed specimens in order to confirm the absence of silver [10].

### 2.2 Antibacterial effect of SBT

The antibacterial activities of SBT specimens on *S. aureus* were quantified using Agar well diffusion technique. 120 μl of *S. aureus* bacteria was spread on Mueller-Hinton Broth (MHB) plates. 4 mm diameter wells sealed with soft agar (0.7%) were taken and the test solution with different concentrations (50, 100 μl) were added into it along with unpolarized and polarized specimens. The plates were then placed for incubation at 37°C in CO_2_ atmosphere for 24 h after which the inhibition zones obtained were measured [19].

### 2.3 Antibacterial rate test

The antibacterial rate test was carried out to quantify the potential of bacteria on the specimens by counting bacteria number for both the polarized and unpolarized specimens taken in a 20 μL (1 × 10^6^ CFU/mL) suspension and placed for incubation at 37 °C for 6 h in CO_2_ atmosphere. Subsequently, the specimens were rinsed with PBS in order to remove the non-adherent bacterial cells. The adherent cells were detached through ultrasonication in 1mL of PBS solution after which 100 μL of bacterial suspension was taken and spread over the agar plate [16]. The number of bacteria on the specimens was counted after incubating the media. The antibacterial ratio was estimated using the equation

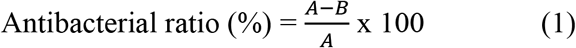

where A is the mean CFUs on control group and B is the mean CFUs on experimental group.

### 2.4 Observation of bacterial adhesion

The evaluation of bacterial adhesion of *S. aureus* was performed by measuring a suspension of 1 × 10^6^ CFU/mL and incubated at 37 °C for 6 h. The specimens were stained with propidium iodide (PI) (81845, Sigma Aldrich) and SYTO 9 (S-34854, Invitrogen™ Thermo Fisher) and the stained specimens were incubated for 15 min. Confocal laser scanning microscopy (CLSM) was employed to qualitatively study the adherent bacteria on the specimens [16].

### 2.5 Bacterial viability test

Gram positive bacteria, *S. aureus* was chosen for the antibacterial study on both types of specimens. *S. aureus* bacterial strain was cultured in MHB medium [20]. The bacterial viability was quantified using MTT assay. The average of three specimens was taken from both the polarized and unpolarized composites. A 24-well plate containing the specimens along with 1 mL *S. aureus* suspension at a concentration of 1 × 10^6^ CFU/mL was incubated at 37 °C in CO_2_ atmosphere. The bacterial culture was carried out for 10, 20 and 30 hours and subsequently *S. aureus* adhered to the culture plates was quantified for bacterial viability after adding 200 μL of bacterial suspension with equal quantity of MTT assay (5 mg/mL) and again incubated for 6 h at 37 °C under CO_2_ atmosphere that resulted in forming formazan crystals. The optical density (OD) values of the specimen solutions was quantified by a spectrophotometric microplate reader at 490 nm [16].

### 2.6 Statistical analysis

All the experimental values were shown as mean ± standard deviation and the differences in data were diverged by one way analysis of variance (ANOVA).

## 3. Results and discussion

The surface of bacterial cells has a heterogeneous complex structure carrying negative charge developed from dissociation of amino-, carboxyl- and phosphate- groups. The charge transporting protein forms pores on the bacterial membranes. The interaction of bacteria with the specimens is dependent on the surface charge and composition of the later. Bacteriostasis in prosthetic bioimplants have been proposed to have developed from the exposure of bioimplants to external electric or magnetic fields [16, 21].

An alternative advanced technique for annihilating the adhesion of bacteria and prevention of biofilm has been highly successful by inducing surface charge on bioimplants. The surface charge can be induced by treating the bioimplant surface with very high strength electric field at higher temperature. It has been observed that polarized bioglass shows antibacterial activity against *S. mutans* [22, 23].

The formation of SBT nanoparticles was confirmed from the XRD pattern that showed aurivilius structure and is depicted in Fig. 1. The size of the SBT nanoparticles were estimated by calculating the full width at half maximum (FWHM) values for (101) plane that indicated the crystallite size to be less than 100 nm.

**Fig. 1.**
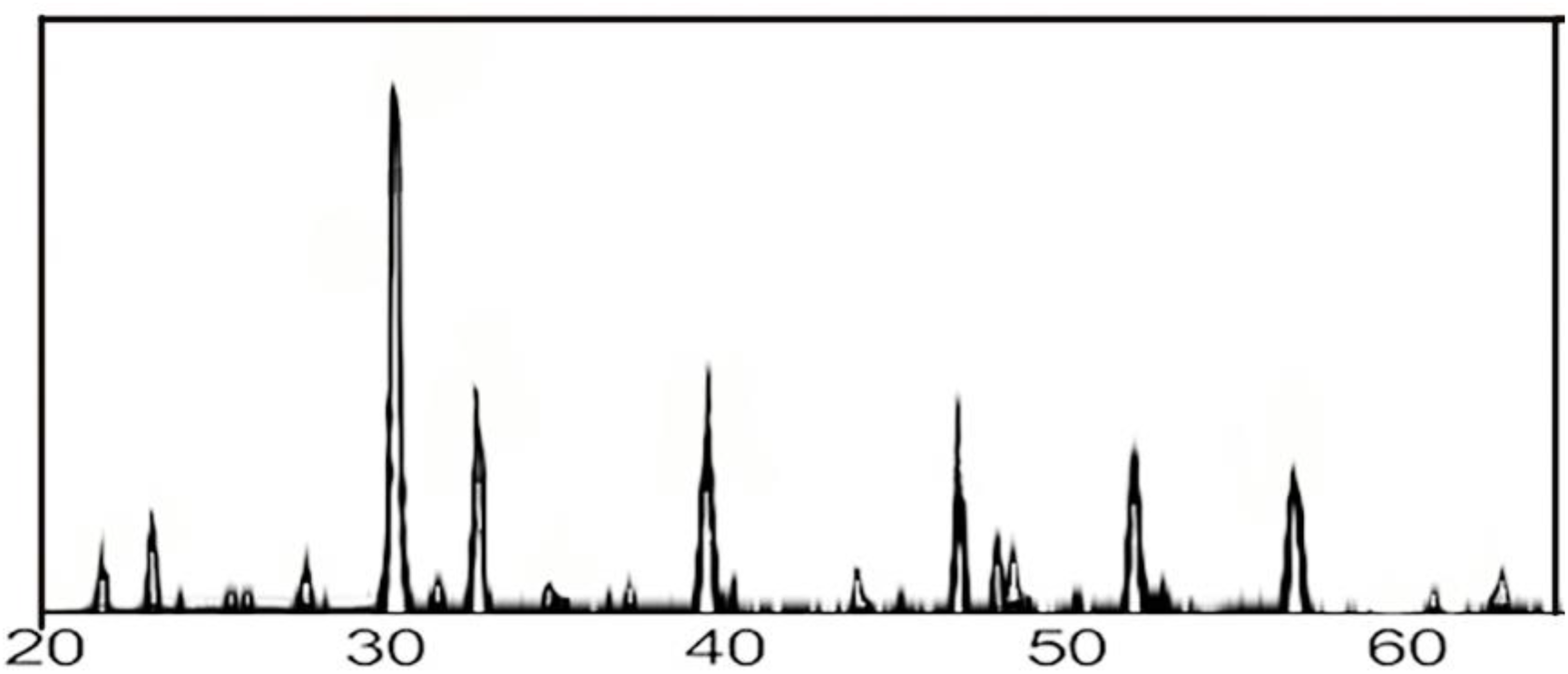
XRD pattern of SBT nanoparticles

The antibacterial activities of SBT NPs were assessed using *S. aureus* that exhibited the size of zones of inhibition (ZOI) to be in the range of 18 mm for unpolarized SBT specimen while 28 mm for the polarized SBT specimens (Table 1). While the unpolarized specimen exhibited lower values of ZOI, the polarized specimen showed to have higher antibacterial activity on *S. aureus* bacteria.

**Table 1:**
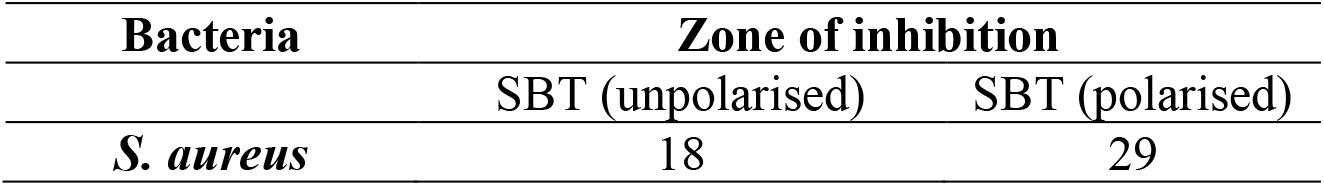
Zone of inhibition as observed on the bacteria against the SBT composites

Studies from literature demonstrate that antibacterial activities may be linked to high voltage pulse applied to bacteria. The primary factor of bactericidal action may be attributed to electroporation which is employed to perforate the bacterial cell membranes. Thus, it’s not mandatory to kill the cells directly while a cell can die once it’s cell membrane is ruptured. Irrespective of the type or size of the cell, perforation by electroporation requires 1 volt or more per cell [24]. Since maximum number of cells have size of approximately 1 μm, the electric field intensity to be applied should be above 1 V·μm^−1^.

Moreover, few other studies have confirmed the cessation of metabolism in bacterial cells under the application of external electric field thus killing bacteria or weakening the activities of bacteria. The bactericidal action of SBT may arise from the presence of antibacterial action arising due to Bismuth (Bi). Bi has been found to have antibacterial action simultaneously providing biocompatibility [25].

The percentage reduction of bacterial cells after 6 h of culture was quantified that showed polarized SBT specimens reducing the bacterial number significantly as compared to the unpolarized specimens. Unpolarized SBT specimens exhibited inadequate killing efficiency for *S. aureus* species. While, the polarized SBT completely killed the bacteria, only 24% bacteria were killed by unpolarized specimens (Fig. 2).

**Fig. 2.**
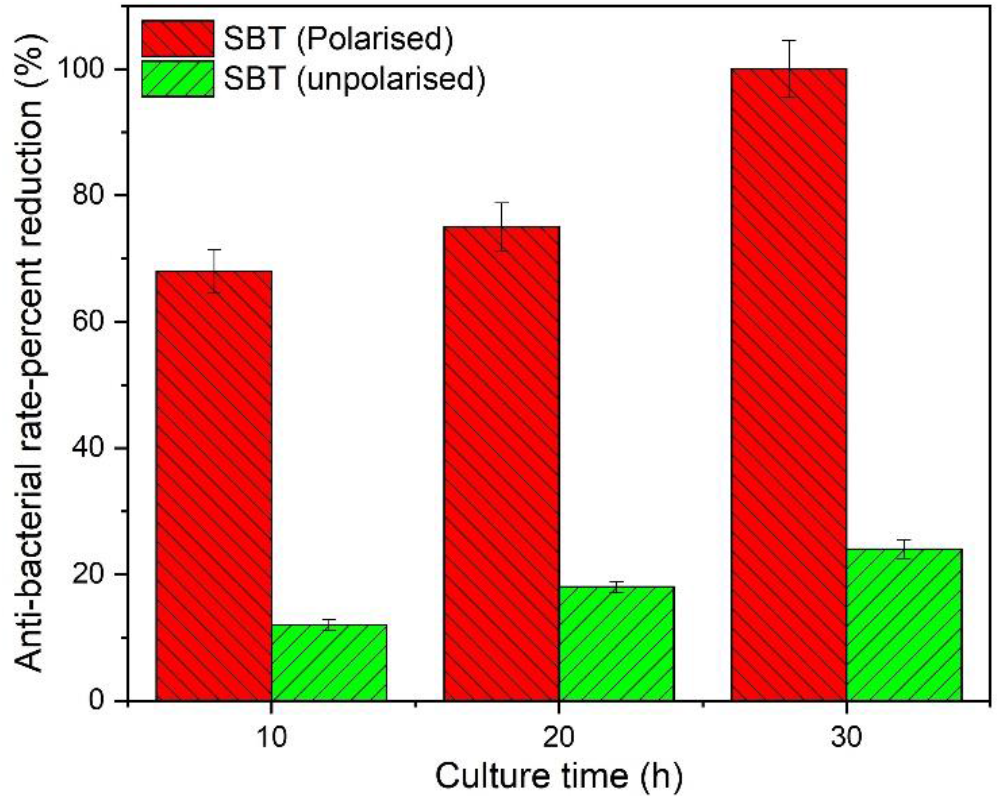
Antibacterial rate- percent reduction of *S. aureus*

The possible adhesion of *S. aureus* on both the polarized and unpolarized SBT specimens was qualitatively characterized by CLSM analysis that depicted live-dead cells. While poor antibacterial activities were witnessed in unpolarized SBT specimen, higher bactericidal action was visible in polarized specimen that shows red spots as dead cells in 6 h of *S. aureus* culture (Fig. 3).

**Fig. 3.**
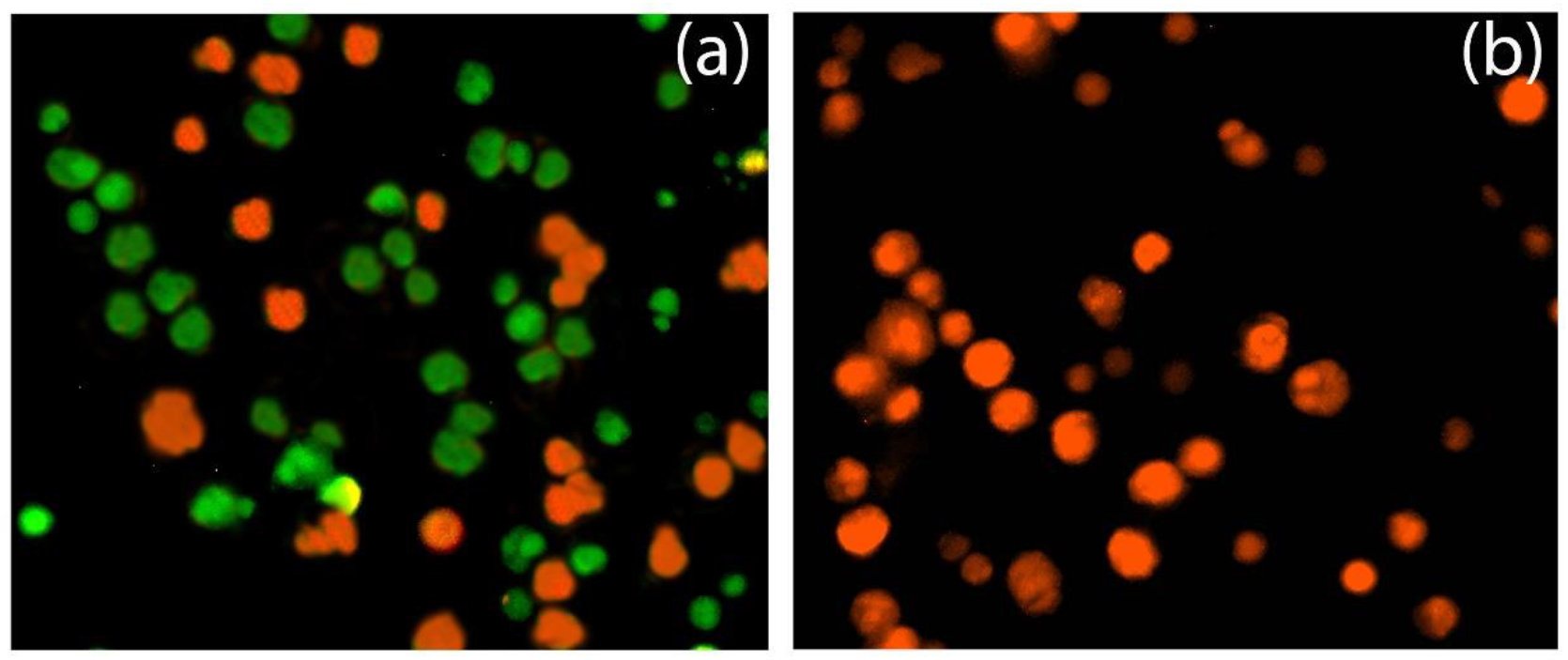
CLSM analysis of *S. aureus* cultured on (a) unpolarized SBT and (b) polarized SBT

After 10, 20 and 30 hours of culture using MTT assay, polarized SBT specimens exhibited restrained growth of bacteria and these results are in consistent with the qualitative CLSM analysis and quantitative ZOI bacteriostasis rate results (Fig. 4).

**Fig. 4.**
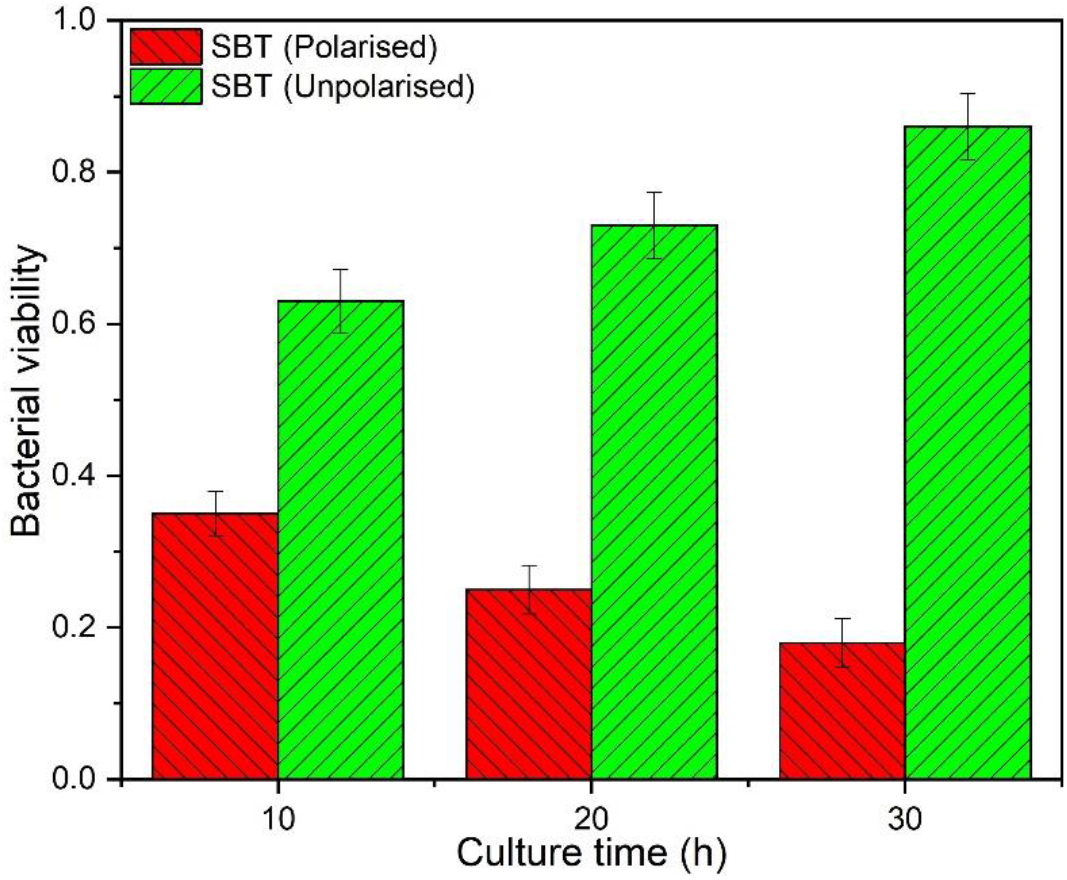
Bacterial viability of *S. aureus* on polarized and unpolarized specimens

*S.aureus* was chosen in this investigation since it’s a very common bacterial species present in chronic and infected wounds and has been studied extensively. Since it’s a Gram +ve bacteria, it is more resistant to electrical stimulation as compared to the Gram −ve bacteria. The higher resistance of Gram +ve bacteria may arise from the thickness/composition of its cell membrane that explains the impact of electrical stimulation producing bacteriostasis effect on Gram +ve bacteria. Hence, *S. aureus* could be taken as an indicator for testing the efficacy of various electrical stimulators.

## 4. Conclusions

The antibacterial effect observed in unpolarized SBT NPs may be ascribed to the presence of Bi which is an antibacterial element. On the other hand, positively polarized SBT specimens had significantly higher bactericidal effect on *S. aureus*. Thus, polarization of a biomaterial can be taken as an advanced method of combating bacterial growth on biomaterials simultaneously preventing the formation of biofilms.

